# A FRET-based high-throughput screening assay for the discovery of *Mycobacterium tuberculosis* DNA ADP-ribosylglycohydrolase DarG inhibitors

**DOI:** 10.1101/2025.07.02.662713

**Authors:** Men Thi Hoai Duong, Yang Lu, Ivan Ahel, Lari Lehtiö

## Abstract

DarTG2 is a conserved toxin-antitoxin ADP-ribosylation system that regulates bacterial survival and the anti-phage response found in many pathogenic bacteria, including *Mycobacterium tuberculosis*. While DNA ADP-ribosyltransferase (DarT2) toxin mono-ADP-ribosylates a single-stranded DNA sequence motif and potentially induces bacterial dormancy, its cognate DNA ADP-ribosylglycohydrolase (DarG) antitoxin reverses the modification and restores bacterial growth. Therefore, developing DarG selective inhibitors may represent a promising and novel strategy to combat drug-resistant tuberculosis. However, no small molecule inhibitors targeting DarG have been identified to date and establishing a high-throughput screening assay based on its DNA ADP-ribosylhydrolase activity would be challenging. Here, we developed and optimized a simple and robust fluorescence resonance energy transfer (FRET)-based high-throughput screening assay to identify small molecule inhibitors targeting DarG macrodomain in *M. tuberculosis*. We generated a FRET pair of *M. tuberculosis* DarG macrodomain and poly-ADP-ribosylated peptide fused with compatible fluorophores. Screening the target-focused phenotypic library using this method led to the identification of pranlukast, which selectively inhibited the DNA ADP-ribosylhydrolase activity of DarG in *M. tuberculosis* and its orthologues without affecting human mono-ADP-ribosyl binders and erasers. Since pranlukast has previously been reported to reduce *M. tuberculosis* burden, further investigation into its action mechanism in this context would be valuable.

## Introduction

ADP-ribosylation is a reversible modification in which ADP-ribosyltransferases transfer an ADP-ribose unit from β-nicotinamide adenine dinucleotide (β-NAD^+^) to target molecules that can be proteins, nucleic acids or antibiotics [1–3]. Typically, the modification is mono-ADP-ribosylation (MAR), but in case of poly(ADP-ribose) polymerases (PARP1, PARP2) that detect DNA damage [4,5] and in case of tankyrases (TNKS1, TNKS2) that control stability of proteins [6], the modification can be extended to branched or linear poly-ADP-ribose (PAR) chains, respectively [7–9]. *In vitro*, PAR chains vary in length and can extend up to 200 ADP-ribose units interconnected by glycosidic bonds, with branching observed approximately every 20 to 50 residues [7]. PAR chains are reported to exhibit a helical secondary structure, showing structural similarities to RNA and DNA [10]. This is evidenced by the cross-reactivity of antibodies raised against PAR with both RNA and DNA [11,12]. ADP-ribosylation plays important roles in many cellular processes, such as DNA damage repair, stress and immune response, chromatin and transcriptional regulation, protein biosynthesis, and cell death [13–15]. Proteins that contain a macrodomain, an evolutionarily conserved protein fold, can bind to or act as ADP-ribosyl hydrolases to degrade this modification [16,17]. Macrodomains are widely identified in all kingdoms of life from virus to human and implied to be involved in a variety of pathogenesis of human diseases such as cancer, neurodegeneration, and bacterial-, viral-mediated infection [17–20].

ADP-ribosylation can be also utilized by toxin-antitoxin (TA) systems which contribute to the pathogenesis of many infectious diseases [21–23]. Recently, the ADP-ribosylation-dependent DarTG2 TA system, identified in various bacteria including the global pathogen *Mycobacterium tuberculosis*, has gained attention due to its significant role in regulating cell growth and inducing bacterial persistence [24–27]. Moreover, this TA system represents the first known example of reversible DNA modification mediated by ADP-ribosylation. In detail, the toxin DNA ADP-ribosyltransferase (DarT2) modifies thymidine bases in a single-stranded chromosomal bacterial DNA. The sequence motif targeted by DarT2 can diverge depending on the species, DarT2 from thermophilic bacterium *Thermus aquaticus* targets TNTC motif while *M. tuberculosis* orthologue prefers TTTT and TTTA motifs [24,25]. As a result, the modification blocks DNA replication, slowing growth of the bacteria [24,25,28]. The antitoxin DNA ADP-ribosylglycohydrolase (DarG) is a homolog of human terminal ADP-ribose glycohydrolase 1 (TARG1) and possesses a conserved N-terminal macrodomain that specifically removes ADP-ribosyl groups from DarT2- modified DNA, thereby preventing excessive toxicity [25,29]. Further investigation showed that partial depletion of DarG triggers a DNA-damage response and sensitizes *M. tuberculosis* to drugs targeting DNA metabolism and respiration [30]. This function makes DarG a crucial component and a potential drug target for tuberculosis and other diseases caused by pathogens that express DarTG2 TA system [26].

The emergence of multidrug-resistant strains of *M. tuberculosis* has posed a serious challenge to tuberculosis control efforts [31–33]. The high rates of antibiotic resistance highlight the urgent need for alternative treatment strategies. Since the DarTG2 system causes bacteriostatic effects, this TA module is recognized as an alternative potential drug target for antibiotic-resistant *M. tuberculosis* [25,26,30]. The potential of targeting ADP-ribosylation as a therapeutic intervention has remained largely unexplored, primarily due to the limited understanding of reversible ADP-ribosylation and its functions in bacterial signaling pathways [20,34,35]. Development of small molecule inhibitors targeting this TA system could provide powerful tools, chemical probes, to gain a better understanding of the DarTG2 system and potentially provide a new avenue for tuberculosis treatment.

Here, we describe the development of an *in vitro* assay based on fluorescence resonance energy transfer (FRET) to screen inhibitors for DarG macrodomain in *M. tuberculosis* (*Mtb*DarG MD). The method uses fluorescent fusion proteins expressed in *E. coli*, resulting in a robust and fast high-throughput assay. The assay measures ratiometric FRET (rFRET) signals upon the binding of mCerulean (CFP) fused *Mtb*DarG MD to the PARylated mCitrine (YFP)-GAP where GAP-tag refers to a C-terminal peptide of a Gαi protein. The assay was validated for high-throughput screening (HTS) in 384-well plate format and tested by screening a target-focused phenotypic library, allowing us to identify pranlukast as the first small molecule inhibitor targeting *Mtb*DarG MD. Pranlukast had an IC_50_ of 4 µM and was selective for *Mtb*DarG MD over a wide panel of human macrodomains. Moreover, pranlukast inhibited the enzymatic activity of *Mtb*DarG and its bacterial orthologues including DarG in *Thermus aquaticus* and SCO6735 in *Streptomyces coelicolor,* suggesting the broad-spectrum inhibition of pranlukast against thymidine-linked DNA ADP-ribosylhydrolases.

## Results

### Set up a FRET assay

We previously showed that TARG1, which is structurally similar to DarG [25], showed a high rFRET signal for the interaction in the FRET pair of CFP-TARG1 and MARylated YFP-GAP [36]. We therefore hypothesized that *Mtb*DarG MD might similarly bind to a MARylated peptide as TARG1 does and tested the applicability of FRET-based assay using CFP-*Mtb*DarG MD and MARylated YFP-GAP. However, no FRET signal indicative of the binding was observed (**Fig. S1A**). Moreover, nanoDSF analysis showed that free ADP-ribose did not stabilize the *Mtb*DarG MD (**Fig. S1B**). These findings suggest that *Mtb*DarG MD requires the DNA attached to the ADP-ribose for efficient binding. The DNA modified by DarT would, however, be hydrolyzed by *Mtb*DarG MD and therefore a mix-and-measure type of high-throughput assay would require more complicated procedures and preparation of specific MARylated and tagged DNA oligonucleotides.

Given that *Mtb*DarG binds to MARylated DNA [25] and PAR chains can mimic the single-stranded DNA (ssDNA) molecule [10,12], we hypothesized that PAR chains might facilitate binding of *Mtb*DarG. Indeed, the FRET-based assay using CFP-*Mtb*DarG MD and PARylated YFP-GAP yielded a detectable rFRET signal, indicating the energy transfer from CFP to YFP. We described here in detail the principle of FRET-based assay for CFP-*Mtb*DarG MD and PARylated YFP-GAP (**Fig. 1A**). The construct of *Mtb*DarG MD was recombinantly expressed as a fusion protein with fluorophore mCerulean CFP. The fluorophore mCitrine YFP was fused with a C- terminal Gαi-based 10-mer peptide (GAP-tag), which was subsequently MARylated at cysteine residue by pertussis toxin subunit S1 (PtxS1) [36]. To extend the PAR chain from the MARylated YFP-GAP, we utilized PARP2, resulting in the formation of the branched PAR chain. Upon excitation of the reaction mixture containing the FRET pair at 410 nm, the rFRET signal was calculated as the ratio of the fluorescence intensity of the YFP emission at 527 nm to the fluorescence intensity of the CFP emission at 477 nm. Binding interactions between the *Mtb*DarG and PAR chain led to a decrease in CFP emission and an increase in YFP emission, thereby enhancing the rFRET signal (**Fig. 1B**). Guanidine HCl (GdnHCl) used as a negative control unfolds proteins and disrupts the binding interaction, but at the used 0.6 M concentration it does not interfere with the fluorescence of CFP and YFP [36].

**Figure 1:**
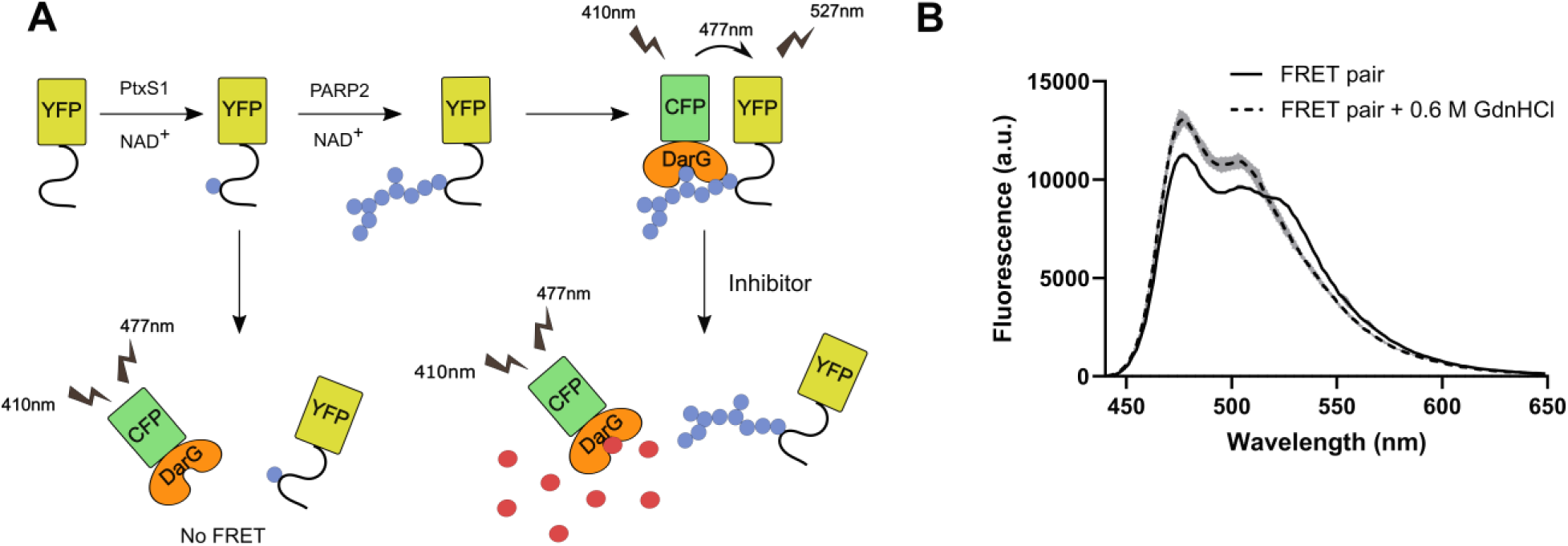
FRET-based assay of CFP-*Mtb*DarG MD and PARylated YFP-GAP. (**A**) The principle of the FRET assay. GAP-tag peptide was fused to YFP and modified at cysteine residue by PtxS1. The PAR chain was extended using PARP2. CFP-fused *Mtb*DarG MD binds to the PAR chain, resulting in FRET signals. Inhibitors can bind to the ADP-ribose binding pocket of *Mtb*DarG MD and disrupt the FRET pair interaction, leading to a loss of FRET signal. (**B**) The fluorescence spectra of 100 nM CFP-*Mtb*DarG MD and 500 nM PARylated YFP-GAP mixture in the absence (solid line) or presence (dashed line) of 0.6 M GdnHCl. The curves shown are mean ± SD of four replicates.

### Signal verification for *Mtb*DarG MD and PAR binding

To confirm the specific interaction between CFP-*Mtb*DarG MD and PARylated YFP-GAP, we investigated whether the removal of the *Mtb*DarG macrodomain and the PARylated GAP from the fluorophores would result in the loss of the FRET signal. Since the expression constructs were designed with a tobacco etch virus (TEV) cleavage site between the *Mtb*DarG macrodomain or PARylated-GAP and the fluorophores, the FRET reaction mixture was treated with 100 nM TEV protease to cleave away CFP and YFP tags. The results demonstrated that the presence of 100 nM TEV led to the decrease of the rFRET signal, which gradually dropped to the basal level after approximately 17 min (**Fig. 2A**). Additionally, we assessed the effect of poly(ADP-ribose) glycohydrolase (PARG), known for its hydrolysis activity against PAR chains but not MARylated substrates [37], on the interaction between *Mtb*DarG MD and PAR. At a tested concentration of 0.5 nM, PARG effectively hydrolyzed PAR chains and rapidly reduced the rFRET signal to the basal level within around 8 min (**Fig. 2B**). These findings implicate a specific interaction between *Mtb*DarG MD and the PAR chain, and that the MARylated GAP-tag does not bind to *Mtb*DarG consistent with our earlier result.

**Figure 2:**
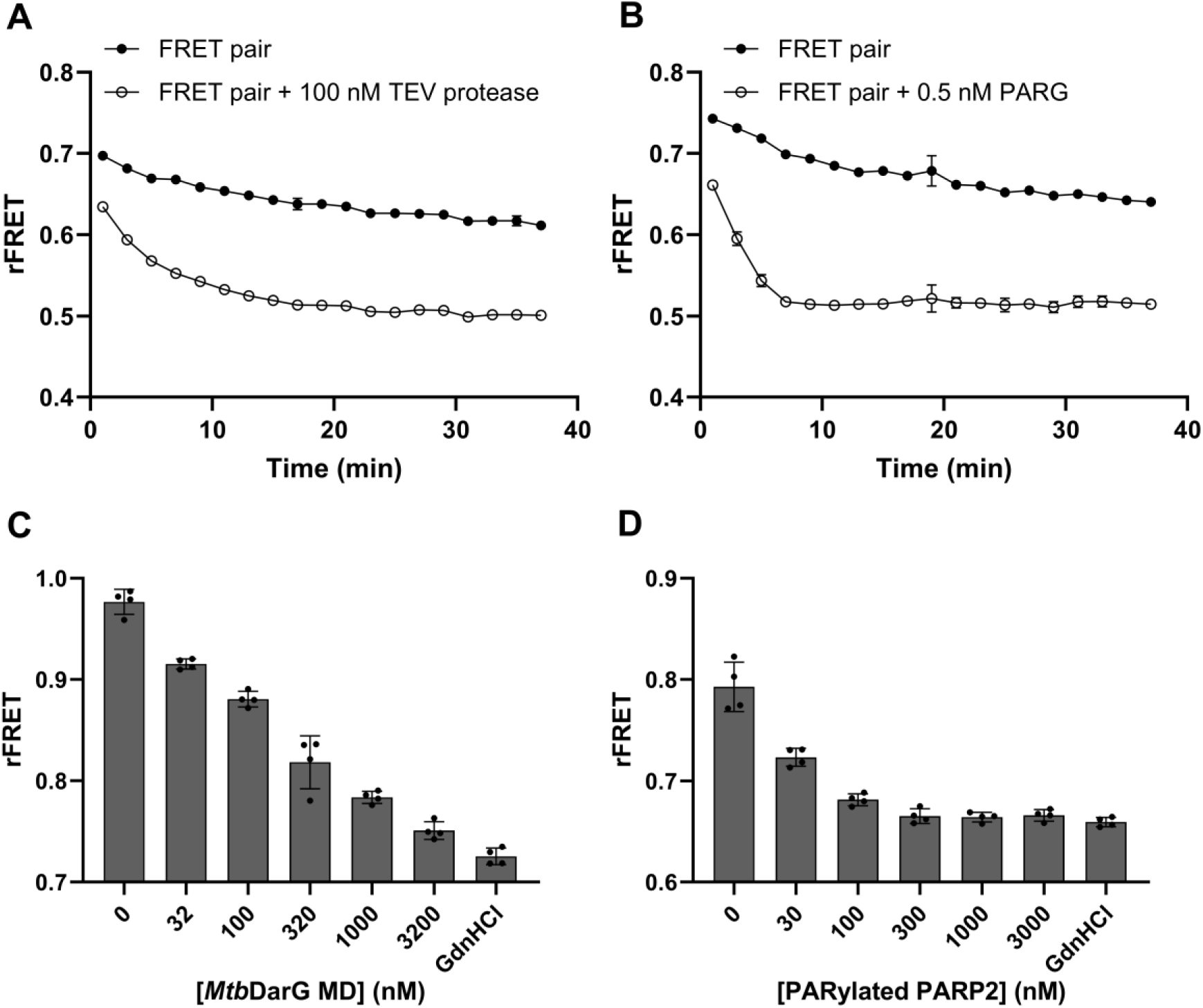
Signal verification of CFP-*Mtb*DarG MD and PARylated YFP-GAP FRET pair. (**A**) Effect of TEV protease on the rFRET signal of *Mtb*DarG MD and PARylated GAP FRET pair. 100 nM CFP-*Mtb*DarG MD was mixed with 500 nM PARylated YFP-GAP in the absence or presence of 100 nM TEV protease. Buffer condition: 10 mM BTP pH 7.0, 0.01% (v/v) Triton X-100, 0.5 mM TCEP, 25mM NaCl. (**B**) Effect of PARG on the rFRET signal of *Mtb*DarG MD and PARylated GAP FRET pair. 100 nM CFP-*Mtb*DarG MD was mixed with 500 nM PARylated YFP-GAP in the absence or presence of 0.5 nM PARG. Buffer condition: 10 mM BTP pH 7.0, 0.01% (v/v) Triton X-100, 0.5 mM TCEP, 25mM NaCl. (**C**) 30 nM CFP-*Mtb*DarG MD and 300 nM PARylated YFP-GAP FRET pair was mixed with increasing amounts of unlabeled *Mtb*DarG in optimized FRET assay buffer: 10 mM MES, pH 6.0, 25 mM NaCl, 0.01% (v/v) Triton X-100, 0.5 mM TCEP, 10% (v/v) glycerol. (**D**) 10 nM CFP-*Mtb*DarG MD and 100 nM PARylated YFP-GAP FRET pair was mixed with increasing amounts of PARylated PARP2 in optimized FRET assay buffer: 10 mM MES, pH 6.0, 25 mM NaCl, 0.01% (v/v) Triton X-100, 0.5 mM TCEP, 10% (v/v) glycerol. Data shown are mean ± standard deviation from four replicates.

However, we noticed that the rFRET signals of samples without TEV protease or PARG treatment were unstable and slightly decreased over the testing time (**Fig. 2A-B**). This instability might be due to the protein aggregation or unstable complex formation in tested buffer consisting of 10 mM BTP pH 7.0, 25 mM NaCl, 0.01% (v/v) Triton X-100, 0.5 mM TCEP. Therefore, we next optimized the buffer condition to gain stable signals before proceeding with further experiments (**Fig. S2-S4**). The optimum pH for the signal was slightly acidic at 6.0 (**Fig. S2**) and was sensitive to increased salt concentration at the low protein concentrations of 500 nM PARylated YFP-GAP and 100 nM CFP-*Mtb*DarG used in the assay (**Fig. S3**). We also observed that in the presence of 5-10% (v/v) glycerol, the signals were significantly more stable over time (**Fig. S4**) and subsequently selected the final buffer conditions as 10 mM MES, pH 6.0, 25 mM NaCl, 10% (v/v) glycerol, 0.01% (v/v) Triton X-100 and 0.5 mM TCEP. This buffer, referred to as FRET assay buffer, was utilized for all subsequent experiments.

To further confirm specific interaction between *Mtb*DarG MD and PAR chain, we conducted a competition assay using unlabeled *Mtb*DarG MD and PARylated PARP2. The results showed that the addition of unlabeled proteins led to a concentration-dependent reduction in the rFRET signal (**Fig. 2C-D**). As both unlabeled *Mtb*DarG MD and additional PAR effectively dropped the rFRET signal, the measured FRET signal is due to a specific binding interaction and not just non-specific aggregation.

### Dissociation constant of *Mtb*DarG macrodomain and PAR chain interaction

To determine the dissociation constant (K_d_) of the interaction between *Mtb*DarG MD and PAR chain, we employed the FRET method as described by Song et al [38]. The FRET emission (EmFRET) was assessed by mixing a constant concentration of CFP-*Mtb*DarG MD (30 nM) with increasing concentrations of PARylated YFP-GAP (0-1.5 µM) to achieve saturation of the CFP with YFP signal. The saturation curve was plotted with EmFRET against the concentration of PARylated YFP-GAP. Since the binding stoichiometry is unknown, the data were fitted using a 1:1 binding model. The resulting saturation curve showed that the apparent binding affinity of *Mtb*DarG MD and PAR was 316 ± 34.7 nM (**Fig. 3A**).

**Figure 3:**
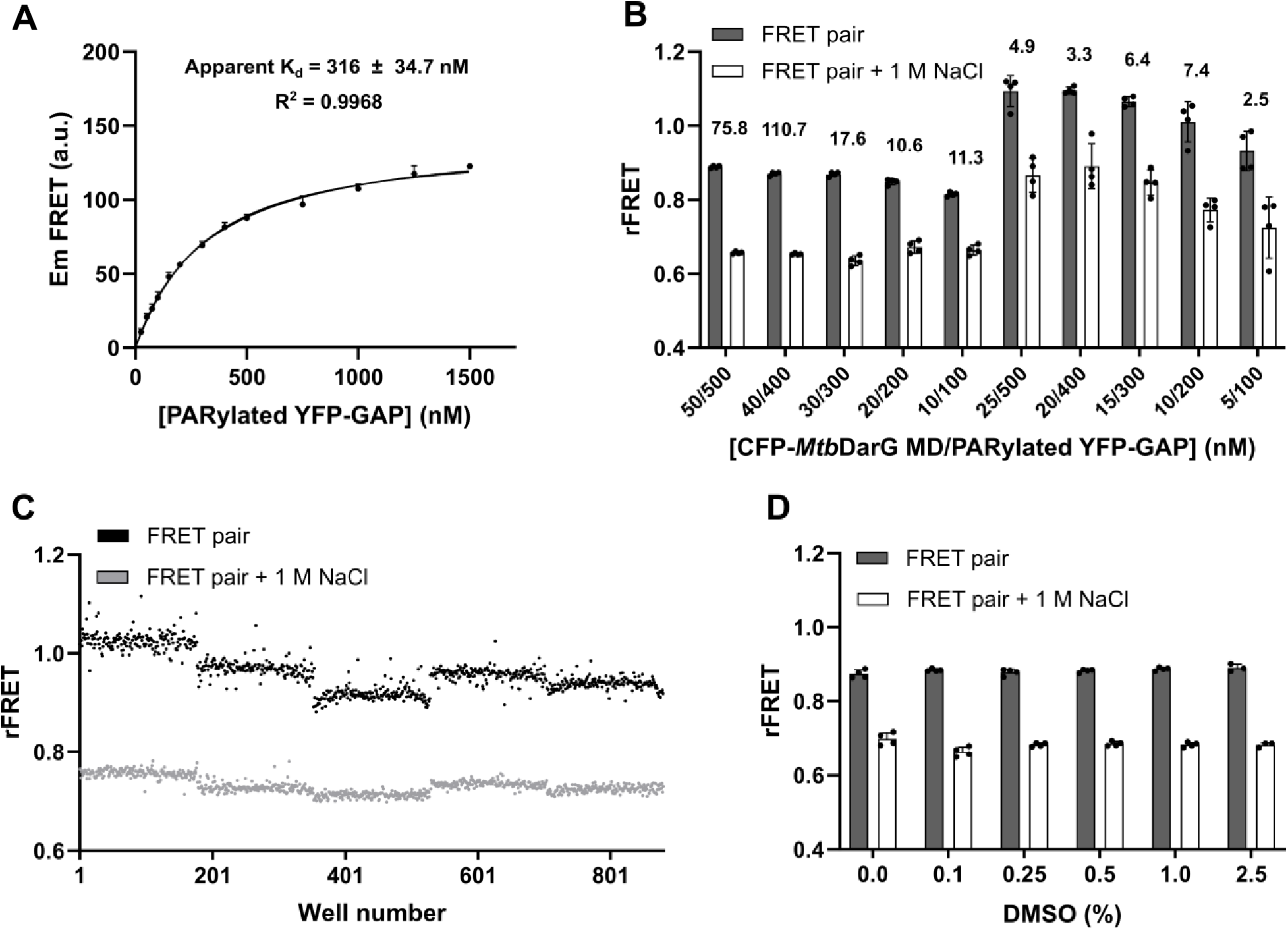
FRET-based assay optimization and signal validation. (**A**) Determination of the apparent FRET-based dissociation constant for CFP-*Mtb*DarG MD and PARylated YFP-GAP. (**B**) Determination of reaction concentrations for CFP-*Mtb*DarG MD and PARylated YFP-GAP. The signal-to-noise ratio is presented on top of each condition. (**C**) Distribution of positive control and negative control signals over the five plates. 30 nM CFP-*Mtb*DarG MD was mixed with 300 nM PARylated YFP-GAP in the absence (positive control) and presence (negative control) of 1 M NaCl. 176 replicates for each positive and negative controls were dispensed in 384-well plate. (**D**) DMSO test indicated that the assay performance was not affected in the presence of DMSO up to 2.5%. Data was analyzed using unpaired t-test (p>0.05). Data shown in panel A, B, D correspond to mean ± SD with number of replicates n = 4. Data shown in panel C correspond to 880 replicates for each positive and negative control.

### Optimization of the assay conditions and signal validation

Next, we optimized protein concentrations for the HTS assay. It was observed that high salt conditions disrupted the binding interaction between CFP-*Mtb*DarG MD and PARylated YFP-GAP (**Fig. S3**). The buffer containing 1 M NaCl can be used as negative control instead of 0.6 M GdnHCl. We tested different concentration ratios of CFP-*Mtb*DarG MD to PARylated YFP-GAP and found that a ratio of 1:10 produced consistent signals with higher signal-to-noise ratios compared to a ratio of 1:20 (**Fig. 3B**). For the HTS assay, we selected conditions with 30 nM CFP-*Mtb*DarG MD and 300 nM PARylated YFP-GAP to minimize protein consumption, making it suitable for large-scale screening campaigns.

After establishing the optimal conditions for the assays, we evaluated the assay quality to ensure its robustness for HTS. The validation process involved assessing the reproducibility of rFRET signals across multiple plates and different days. We used 384-well plates, with each plate containing 176 replicates of the negative control (FRET pair in the presence of 1 M NaCl) and 176 replicates of the positive control (FRET pair only) and measured the rFRET signals (**Fig. 3C**). Statistical parameters demonstrating the assay quality for HTS are summarized in **Table S1**. The Zʹ-factors, which measure the separation between the negative and positive control signals, were calculated for five different plates and found to be between 0.67 and 0.75. The Z’-factor above 0.5 indicates that the FRET signal shows sufficient separation between negative and positive control and the assay is considered excellent for HTS [39]. Our results confirm that the FRET assay is robust, reproducible, and suitable for reliable HTS.

Small molecule compounds in screening libraries are typically dissolved in DMSO, which can potentially impact assay performance. To ensure the robustness of our assay, we evaluated the effect of DMSO, up to 2.5%, on the rFRET signal. The results demonstrated that the assay performed well in the presence of DMSO at least up to 2.5% (v/v), indicating that the assay is robust and suitable for compound screening under these conditions (**Fig. 3D**).

### Validatory screening

To validate the assay for inhibitor screening, we applied this FRET-based assay to screen the target-focused phenotypic screening library to identify the inhibitors for *Mtb*DarG MD and PAR interaction. This library consists of 1832 bioactive compounds with established mechanisms covering more than 600 drug targets. Therefore, the potential hit compounds could be suitable to be tested in the *in vivo* system directly to validate the targeting of *Mtb*DarG. Compounds were screened at a single concentration of 20 µM, and reaction volume was 10 μL. At this concentration, the assay contained 0.02% (v/v) DMSO which was shown to have no effect on the assay performance (**Fig. 3D**).

For the screening, we implemented several approaches to exclude compounds that non-specifically inhibit the FRET interaction or exhibit intrinsic fluorescence at the wavelengths measured. Intrinsic fluorescence of certain compounds can interfere with the ratiometric signal, leading to false positives or inaccurate readings. The exclusion process is summarized in **Fig. 4A**. Firstly, compounds were excluded if their raw fluorescence values were 120% higher than that of the positive control at the 527 nm readout or 80% lower than that of the positive control at the 477 nm readout. These thresholds were chosen to identify and remove compounds that exhibited excessive intrinsic fluorescence, which could interfere with accurate FRET signal measurements. This initial filter successfully excluded 154 compounds (8.4%). Secondly, we applied an additional excitation wavelength of 430 nm for FRET measurement to further identify compounds with significant fluorescence interference. Compounds with such interference typically exhibit stronger excitation at either 410 nm or 430 nm, resulting in discrepancies in the rFRET signals between these two excitation wavelengths. Compounds showing more than a 30% difference in apparent inhibition between the 410 nm and 430 nm readouts were excluded, eliminating an additional 59 compounds (3.2%).

**Figure 4:**
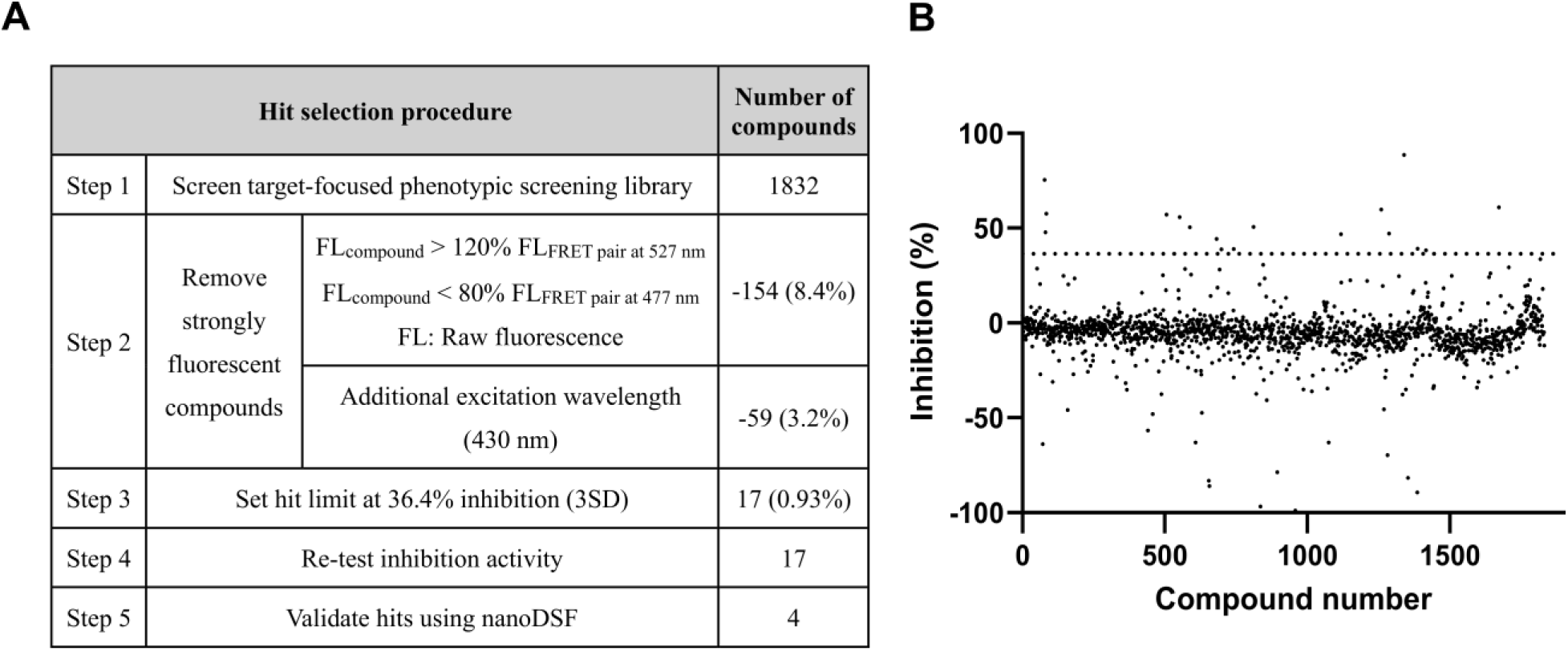
Screening of the target-focused phenotypic screening library for *Mtb*DarG MD and PARylated GAP FRET pair. (**A**) Summary of hit selection procedure. (**B**) Screening of 1,832 compounds identified several compounds inhibiting the FRET pair binding interaction. The data represent the percentage of inhibition, measured with single readings. Seventeen compounds (hit rate 0.93%) were identified as hits. The dotted line indicates the threshold for hit identification, set at 36.4% (3 x SD above the mean value).

After passing through several filters, compounds with inhibition higher than 36.4% (Three standard deviations above the mean value) were considered hits (**Fig. 4B**). In the primary screening, we identified 17 compounds as initial hits for *Mtb*DarG MD, corresponding to a hit rate of 0.93%. Since the screening was conducted in singlet, we re-tested these hits at 20 μM of compound concentration using the FRET assay to confirm their inhibitory activity. Upon re-testing, all compounds showed the inhibition activity against the *Mtb*DarG MD-PAR interaction, confirming their status as hits (**Fig. S5**).

### Hit validation

To rule out the possibility that the observed rFRET signal disruption is due to protein destabilization, we conducted nano differential scanning fluorimetry (nanoDSF) analysis for identified hit compounds. Our results demonstrated that four compounds, including pranlukast, GW501516, tiplaxtinin, and BMS303141, caused right shifts in the melting temperature (T_m_) of the *Mtb*DarG MD, with increases of 1.18 ± 0.05 °C, 1.52 ± 0.08 °C, 1.31 ± 0.06 °C, and 1.21 ± 0.10 °C, respectively. These shifts suggest a high likelihood that compounds directly bind to and stabilize *Mtb*DarG MD (**Fig. 5A-B**). While pranlukast and tiplaxtinin demonstrated the most potential inhibitory activity, with half-maximal inhibitory concentration (IC_50_) values of 4 µM (pIC_50_ ± SEM: -5.40 ± 0.02) and 5.7 µM (pIC_50_ ± SEM: -5.24 ± 0.03), respectively, the IC_50_ values for GW501516 and BMS303141 were 56.5 µM (pIC_50_ ± SEM: -4.25 ± 0.01) and 18.7 µM (pIC_50_ ± SEM: -4.73 ± 0.03), respectively (**Fig. 5C-F**). However, the IC50 curves exhibited considerable steepness, which may indicate a promiscuous aggregator effect of these compounds or suggest that compounds may interact with multiple distinct sites on the protein [40]. Because 0.01% Triton X- 100 was included in buffer condition to mitigate nonspecific inhibitory effects from compound aggregation, the observed inhibitory activity is less likely due to aggregate formation. In addition, dynamic light scattering data from nanoDSF showed that the presence of the compounds did not increase light scattering relative to the protein-only control, suggesting the absence of detectable aggregates under the tested buffer condition (**Fig. S6**). Nevertheless, nonspecific inhibition by aggregators is a common artifact in HTS and it is important to carefully further validate specificity of these hits before drawing conclusions about their activities.

**Figure 5:**
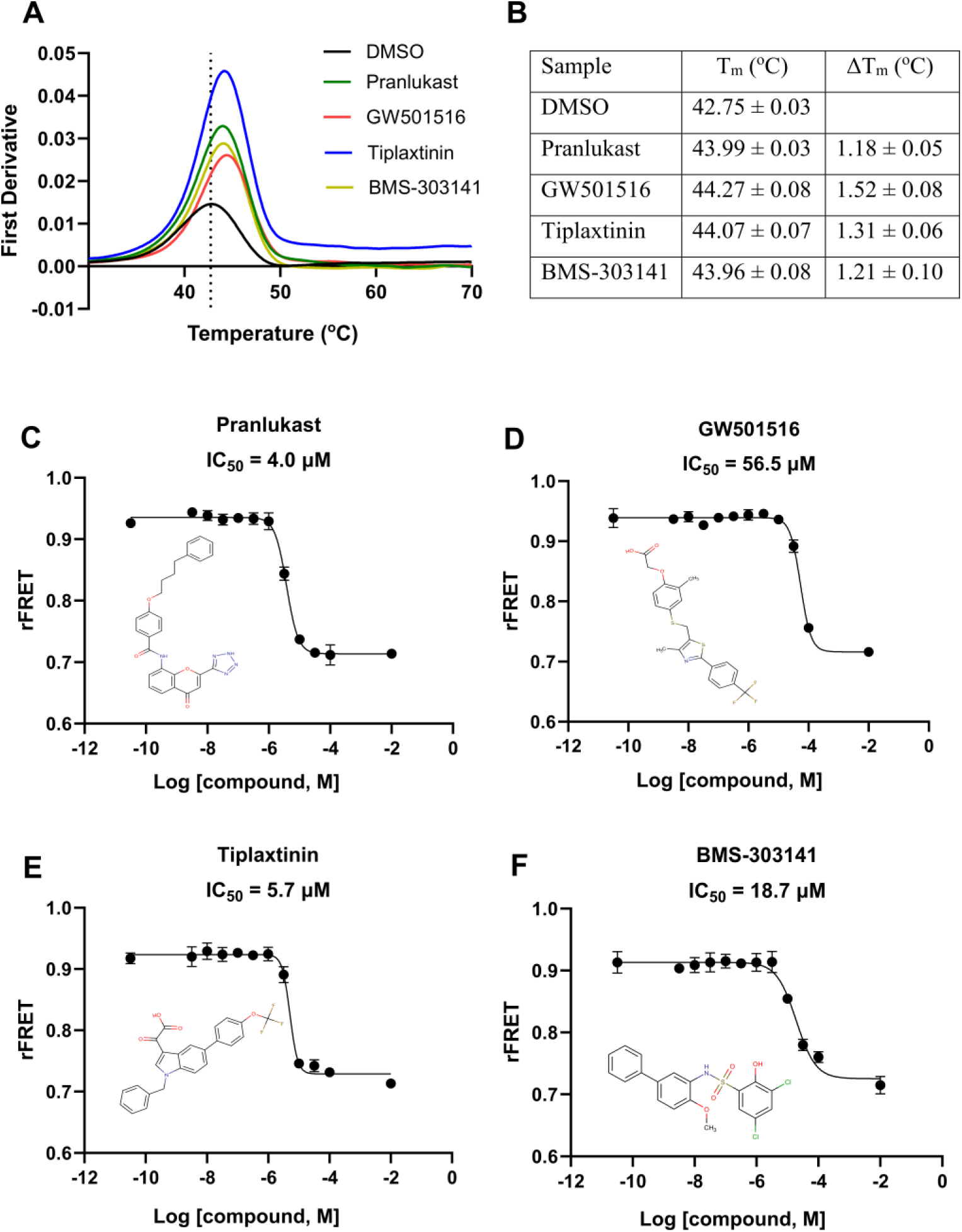
Validation of hit compounds. (**A**) Melting curves of *Mtb*DarG MD were measured using nanoDSF in the absence and presence of 60 µM each hit compound. The experiments were performed in triplicates, and representative curves for the control and each compound are displayed. The dotted line indicates the T_m_ of DMSO control sample. (**B**) Summary of *Mtb*DarG MD T_m_ with and without compounds and corresponding T_m_ shifts. Thermal stability increased in the presence of compounds, indicating the binding interaction with the *Mtb*DarG MD. Data shown are mean ± SD with number of replicates n = 3. (**C-F**) IC_50_ curves and chemical structures of four hit compounds. Data shown are mean ± SD with number of technical replicates n = 4. IC_50_ values are displayed as mean ± SD from three independent measurements.

To eliminate the possibility of nonspecific inhibition, we assessed the potential of the compounds to interfere with the binding interactions of various macrodomains. Four compounds were tested at a single concentration of 20 μM using the FRET-based assay, as described earlier [36]. The results indicated that tiplaxtinin and BMS303141 interfered with the binding activity of several macrodomains, including PARP9 MD1, PARP14 MD1, PARP14 MD3, and PARP15 MD2. In contrast, pranlukast and GW501516 had very low or no effect on the binding interactions of other macrodomains, demonstrating their high selectivity against *Mtb*DarG without broadly affecting other macrodomains (**Fig. 6A**). Next, we tested a wider panel of human ADP-ribosyl binders/hydrolases with the most potent inhibitor, pranlukast, to evaluate its potential as an antibacterial agent. Notably, pranlukast remained selective for *Mtb*DarG over a profiling panel including its structurally similar paralog TARG1 even at a high tested concentration of 100 µM. Only weak inhibition was observed for PARP9 MD1, PARP14 MD1 and PARP15 MD1 with IC_50_ values of approximately 100 μM (**Fig. 6B**). While pranlukast was found to inhibit PARG, it also destabilized the protein when measured with nanoDSF.

**Figure 6:**
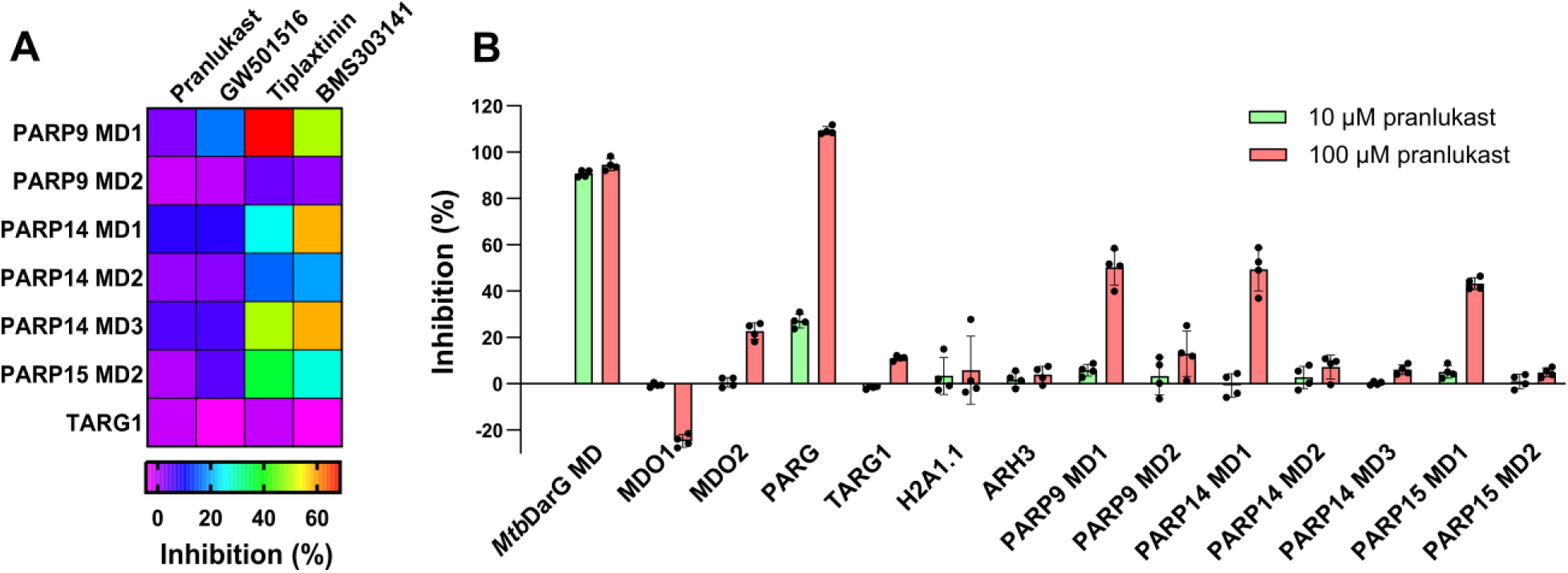
The selectivity of four hit compounds (**A**) Heat map visualizing the inhibition percentages of four hit compounds against a panel of macrodomains. Compounds were tested at a single concentration of 20 µM. The color scale represents the inhibition percentage. The experiments were conducted in duplicate, and the data presented are the mean values derived from duplication. (**B**) A profiling panel of pranlukast against human ADP-ribosyl binders/hydrolases. Data shown are mean ± SD with number of replicates n = 4.

### Effect of pranlukast on hydrolytic activity of DarG

Next, we tested the inhibitory effect of pranlukast on the enzymatic activity of *Mtb*DarG. For the substrate, we utilised a ssDNA oligonucleotide containing a TCTC motif that is specifically ADP-ribosylated by the *Thermus aquaticus* DarT2 toxin at a thymidine residue [25]. Notably, pranlukast inhibited the enzymatic activity of *Mtb*DarG on this ADP-ribosylated ssDNA substrate (**Fig. 7**). In addition, pranlukast exhibited an even higher inhibitory potency against DarG from *Thermus aquaticus* bacterium (*T.aq.* DarG) and SCO6735. SCO6735 is a DNA damage-inducible macrodomain protein from *Streptomyces coelicolor* bacteria, related to DarG and TARG1, that also has the ability to hydrolase ADP-ribosylation from thymine bases [41,42]. These results suggest that pranlukast may have broad-spectrum potential to inhibit bacterial thymidine-linked DNA ADP-ribosylhydrolases.

**Figure 7:**
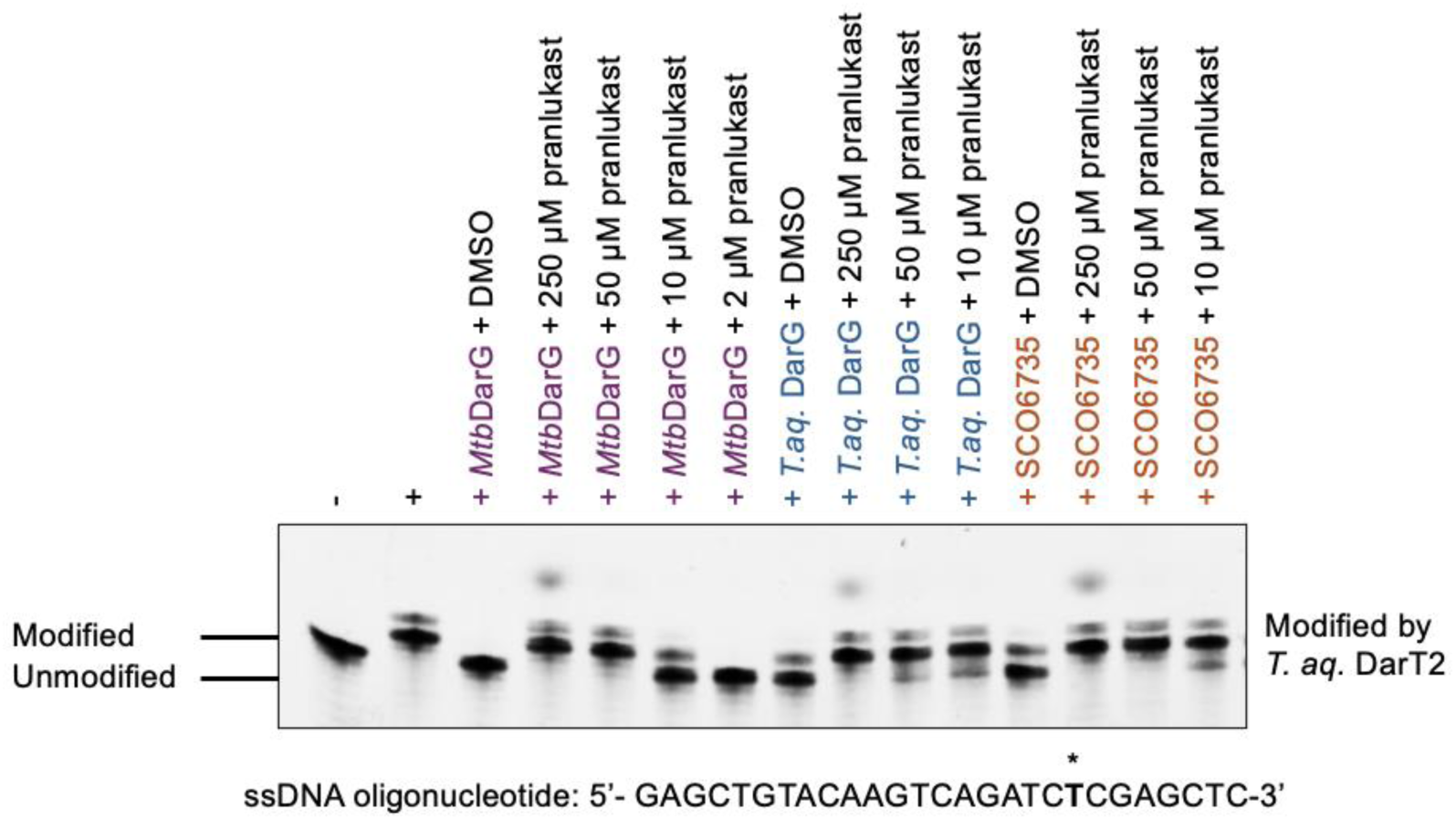
Comparison of pranlukast efficiencies on the enzymatic activities of *Mtb*DarG, *T. aq.* DarG, and SCO6735 in hydrolysing thymidine base ADP-ribosylation catalysed by *T. aq.* DarT2 through pranlukast titrating. *In vitro* ADP-ribosylation assay was conducted using a pre-established ssDNA substrate [24,25], shown above the gel image, with the thymidine base targeted for ADP-ribosylation by *T. aq.* DarT2 indicated by an asterisk.

## Discussion

1. *M. tuberculosis* is the causative agent of tuberculosis, one of the deadliest infectious diseases worldwide. A major challenge in treating tuberculosis is drug resistance, which is largely due to *M. tuberculosis*’s ability to replicate slowly and enter a dormant state. Therefore, understanding and targeting the mechanisms that regulate bacterial replication may provide novel therapeutic strategies. One such mechanism is the DarTG2 TA system, which has emerged as a regulatory checkpoint for controlling bacterial DNA replication [24,28] and disrupting this TA system by small molecule inhibitors may sensitize *M. tuberculosis* to antibiotic treatments. Specifically, the DarG is an essential gene in the context of active DarT toxin protein so targeting the antitoxin component DarG using small molecule inhibitors may represent a promising strategy to kill *M. tuberculosis*.

While HTS assays have been developed to identify inhibitors of various macrodomains [43], the specific function of DarG to target MARylated ssDNA, but not MARylated proteins, presents a unique challenge in design of effective screening assays. Here, we developed a FRET-based assay consisting of a FRET pair of CFP-*Mtb*DarG MD and PARylated YFP-GAP to identify small molecule inhibitors targeting *Mtb*DarG MD. This "mix-and-measure" format eliminates the need for separation steps or specialized reagents and is readily scalable to a standard 384-well plate format, making it suitable for HTS applications. The assay was validated through Z’-factor analysis, demonstrating its robustness and reproducibility. Although it was developed specifically for *Mtb*DarG, the assay can potentially be adapted to screen inhibitors of other DNA-linked ADP-ribosyl binding proteins.

However, the assay has certain limitations. The PARylated substrate used does not accurately reflect the physiologically relevant MARylated ssDNA that DarG interacts with *in vivo*. Additionally, the molecular basis of the interaction between *Mtb*DarG and PAR remains unclear. We hypothesize that *Mtb*DarG possibly binds to PAR chains through the terminal ADP-ribose unit. Electrostatic interactions may also play a role, because the negatively charged diphosphate of PAR could mimic the charge distribution of DNA. We used PARP2 to extend the branched PAR chains, and the effect of linear PAR chains on binding was not evaluated.

Through a validatory screening campaign, the most potent compound identified to inhibit *Mtb*DarG was pranlukast, a selective leukotriene LTC4/D4/E4 receptor antagonist that is FDA-approved for the treatment of bronchial asthma [44]. Notably, pranlukast has previously been reported to act as a novel allosteric inhibitor of *M. tuberculosis* ornithine acetyltransferase, an enzyme involved in arginine biosynthesis, with a dissociation constant of 139 µM [45]. Pranlukast was also shown to significantly reduce bacterial load and granuloma formation in *M. tuberculosis*- infected mouse lungs [45,46]. In our study, pranlukast was found to inhibit the enzymatic activity of *Mtb*DarG, as well as its orthologues including *Thermus aquaticus* DarG and a more diverged protein related to DarG - SCO6735 from *Streptomyces coelicolor*. Notably, its potency against *Mtb*DarG in both the FRET-based binding assay and the hydrolysis assay was higher than that observed for ornithine acetyltransferase. These results justify further investigations to evaluate the mechanism and therapeutic potential of pranlukast against *M. tuberculosis* and other bacterial pathogens.

## Materials and Methods

### Cloning, protein expression and purification

For CFP-*Mtb*DarG MD construct, gene encoding for *Mtb*DarG MD (M1-A155) (Uniprot ID: O53605) was amplified by PCR and then cloned into pNIC-CFP (Addgene #173074). SLIC method was employed for cloning experiments [47]. Briefly, pNIC-CFP was linearized in PCR using site specific primers. 100 ng of the linearized plasmid was mixed with the PCR product in a 1:3 molar ratio. The mixture was then incubated with T4 polymerase for 2.5 min at room temperature, followed by 10 min on ice. After that, the mixture was transformed into *E. coli* NEB 5α competent cells. Colonies were grown on LB agar plate containing selection antibiotic marker, kanamycin, and 5% sucrose as SacB-based negative selection marker [48,49] at 37 °C overnight. The CFP-*Mtb*DarG construct contained *Mtb*DarG macrodomain with an N-terminal 6x His-tag followed by CFP and a TEV protease cleavage site.

The YFP-GAP construct - GAP-tag refers to a C-terminal peptide K345-F354 of Gαi - is available from our previous work (Addgene #173080) [36]. MARylated and PARylated YFP-GAP and proteins used in the profiling were produced as described earlier [36].

CFP-*Mtb*DarG MD plasmid was transformed to *E. coli* BL21 (DE3) competent cells. Transformed cells were inoculated in 5 mL of the Luria Broth media containing 50 μg/mL kanamycin as a selection marker. The pre-cultures were allowed to grow overnight in 37 °C incubator at 200 rpm. 1 L of Terrific Broth autoinduction media including trace elements containing 8 g/L glycerol and 50 μg/mL were inoculated with 5 mL of pre-culture. Flasks were incubated at 37 °C shaker incubator at 220 rpm until the optical density at 600 nm (OD_600_) reached 1. The temperature was reduced to 18 °C and cultures continued growing overnight. The cells were harvested the next day by centrifugation at 4,200 x g for 45 min at 4 °C. The pellets were resuspended in lysis buffer containing 50 mM HEPES, pH 7.5, 500 mM NaCl, 10 mM Imidazole, 10% (v/v) glycerol, 0.5 mM TCEP and stored at -20 °C. CFP-*Mtb*DarG was purified by immobilized metal affinity chromatography (IMAC) and size-exclusion chromatography (SEC). Cell suspensions were thawed and supplemented with 0.1 mg/mL lysozyme, 0.02 mg/mL DNAase, 0.1 mM protease inhibitor pefabloc (Sigma-Aldrich, USA). Cells were then lysed by sonication and the lysate was then centrifuged at 16,000 rpm for 45 min at 4 °C. The supernatant was filtered through 0.45 µm filter before loaded onto a 5 mL IMAC column charged with Ni^2+^ and pre-equilibrated with lysis buffer. The column was then washed with 5 column volumes of lysis buffer followed by another washing with wash buffer containing 25 mM Imidazole. Proteins were eluted by using an elution buffer containing 300 mM Imidazole. Eluted proteins were concentrated using 30 kDa molecular weight cut off (MWCO) concentrator and loaded onto a Hiload^TM^ 16/600 superdex 75 (GE Healthcare, Sweden) gel filtration column which was pre-equilibrated with SEC buffer containing 30 mM HEPES, pH 7.5, 300 mM NaCl, 10% (v/v) glycerol, 0.5 mM TCEP. The purified protein was concentrated, divided into 50 μL aliquots and flash frozen in liquid nitrogen to be stored at - 70 °C. For untagged *Mtb*DarG, an additional step of TEV protease treatment in 1:30 molar ratio was performed, followed by reverse IMAC to remove any impurities from desired protein.

### FRET measurement

The fluorescent signal was measured using Spark or Infinite M1000 pro plate reader (Tecan, Männedorf, Switzerland) and proxiplate-384 F plus (Revvity, Waltham, MA, USA). CFP-fused *Mtb*DarG macrodomain was excited at 410 nm wavelength and fluorescence emission at 477 nm and 527 nm was recorded. Excitation wavelengths were set to 20 nm bandwidth, while emission wavelengths were set to 10 nm bandwidth. The device flashed the light 50 times at a frequency of 400 Hz. Each measurement had an integration time of 20 µs and a settle time of 10 ms.The rFRET value was then calculated by dividing the fluorescence intensity at 527 nm by 477 nm after blank subtraction. An additional excitation wavelength of 430 nm was used in the screening campaign. The experiments were performed in the buffer containing 10 mM MES, pH 6.0, 25 mM NaCl, 10% (v/v) glycerol, 0.01% (v/v) Triton X-100, 0.5 mM TCEP with 10 µL volume per well. This buffer recipe is called FRET buffer for short.

### Competition assay

25 nM CFP-*Mtb*DarG MD was mixed with 300 nM PARylated YFP-GAP with increasing concentrations of unlabelled *Mtb*DarG MD. The FRET pair of 10 nM CFP-*Mtb*DarG MD and 100 nM PARylated YFP-GAP was mixed with increasing concentrations of PARylated PARP2. 50 μM PARylated PARP2 was prepared by incubating a reaction mixture of 50 µM PARP2, 50 μM Oligo and 1 mM NAD^+^ in 30 min at RT as described earlier [36].

### Dissociation constant measurement

A constant concentration of 30 nM CFP-*Mtb*DarG MD was mixed with variable concentrations of PARylated YFP-GAP. Briefly, 6 µM CFP-*Mtb*DarG MD prepared in buffer containing 20 mM HEPES, 150 mM NaCl, 10% (v/v) glycerol, 0.5 mM TCEP, pH 7.5 was transferred to 384-well plate to get the final protein concentration of 30 nM. 50 µM of PARylated YFP-GAP stock was prepared in FRET assay buffer and transferred to 384-well plate to get final PARylated YFP-GAP concentrations of 0, 25, 50, 75, 100, 150, 200, 300, 400, 500, 750, 1000, 1250, 1500 nM. The transfer was done using Echo 650 acoustic liquid dispenser (Beckman Coulter Life Sciences, Indianapolis, IN, USA). Then, the sample wells were backfilled with buffer to 10 μL per well using Mantis liquid handler (Formulatrix, Bedford, MA, USA). The fluorescence was measured using Spark plate reader (Tecan, Männedorf, Switzerland). Upon excitation at 430 nm, the fluorescence emission at 477 nm (10 nm bandwidth) and 527 nm (10 nm bandwidth) were recorded. In addition, the emission at 527 nm (10 nm bandwidth) was also measured after the excitation of 477 nm light (10 nm bandwidth). The analysis was done in GraphPad Prism 10 as described by Song et al. [38].

### FRET assay validation

Assay validation was performed to confirm the assay robustness. A 384-well plate consisted of 176 maximal (positive controls) and 176 minimal (negative controls) signal points and 32 blank wells. Blank wells were filled with only FRET buffer. Maximal signal wells had the FRET pair (30 nM CFP-*Mtb*DarG MD and 300 nM PARylated YFP-GAP) and minimal signal wells had the same FRET pair and 1 M NaCl. Reaction volume was 10 μL per well. In total, five control plates were prepared to test the repeatability between different wells, plates, and days. One plate was measured on the first day, another plate on the second day, and three plates on the third day. Parameters including signal-to-noise ratio (S/N), signal-to-background (S/B), and screening window coefficient Z’ were calculated to access the assay performance [39,50].

### Screening in-house library

The target-focused phenotypic screening library including 10 mM compound stocks dissolved in DMSO was prepared in 384-well polypropylene Echo source plates. 20 nL of compound stocks were transferred to proxiplate-384 F plus reaction plate using Echo acoustic liquid dispenser 650 (Beckman Coulter Life Sciences, Indianapolis, IN, USA). The FRET pair reaction mixture consisted of 30 nM CFP-*Mtb*DarG MD and 300 nM PARylated YFP-GAP. 10 μL of FRET pair reaction mixture per well were dispensed to reaction plates using Mantis liquid handler (Formulatrix, Bedford, MA, USA). rFRET signals were measured and calculated as described above.

### Potency measurements

IC_50_ values were measured to validate hit activity. Serial half-log dilutions of hit compounds were tested, ranging from 100 µM to 0.003 µM. Final DMSO concentration in reaction wells was 1%. The reaction volume was 10 µL per well. FRET pair mixture of 30 nM CFP-*Mtb*DarG MD and 300 nM PARylated YFP-GAP in the FRET buffer was dispensed to reaction wells. Samples in the absence or presence of 1 M NaCl were used as positive or negative controls, respectively. Analysis of data was done using GraphPad Prism version 10 using nonlinear regression analysis.

### Differential scanning fluorimetry

NanoDSF was used to study protein-ligand binding. 0.3 mg/ml *Mtb*DarG MD was mixed with 60 μM of hits in the buffer containing 10 mM MES, pH 6.0, 100 mM NaCl, 5% (v/v) glycerol, 0.5 mM TCEP. Final DMSO concentration in samples was 1.2%. Measurements were performed using Prometheus NT.48 (nanoTemper, Germany). The melting curves were recorded in 1 min intervals from 20–90 °C, with a 1°C increment per min.

### Selectivity measurements

For profiling, 10 nL or 100 nL of 10 mM pranlukast stock was transferred to the reaction plate using Echo 650 acoustic liquid dispenser (Beckman Coulter Life Sciences, Indianapolis, IN, USA). To prepare the FRET pairs, 1 µM of CFP-MAR binder/eraser mixed with 5 µM MARylated YFP- GAP in the assay buffer as described earlier [36]. 200 µM ADP-ribose was used as a negative control. Reaction volume was 10 µL. Final DMSO concentration was 1%.

### *In vitro* ADP-ribosylation activity assays

Proteins used in this assay were expressed and purified as previously published [24,25,42]. Hydrolysis assay was done as before [25,35]. Briefly, ADP-ribosylation reactions were conducted in a reaction buffer containing 50 mM Tris-HCl (pH 7.5), 50 mM NaCl, and 1 mM DTT, with 0.5 µM *T. aq.* DarT2 protein, 3 µM pre-established ssDNA oligonucleotide (5’- GAGCTGTACAAGTCAGATCTCGAGCTC-3’) [24,25], and excess β-NAD⁺ (1 mM), incubated at 37 °C for 30 min. Reactions were terminated by heating the sample at 95 °C for 5 min. Pranlukast was titrated at concentrations of 250, 50, 10, and 2 µM in DMSO, with DMSO alone serving as a control, and pre-incubated with 1 µM of the respective ADP-ribosylhydrolases (*Mtb*DarG, *T. aq.* DarG, or SCO6735) at room temperature for 10 min. De-modification assays were subsequently conducted with the indicated hydrolases in the presence of either pranlukast or DMSO at 37°C for 30 min, after which reactions were terminated by adding urea loading dye (4 M urea, 10 mM Tris-HCl (pH 7.5) and 10 mM EDTA) and heating at 95 °C for 5 min. Final reaction products were separated and analysed on denaturing urea-polyacrylamide gels in 1x Tris/Borate/EDTA (TBE) buffer, stained with SYBR Gold Nucleic Acid Gel Stain (Invitrogen), and visualised under 340-nm ultraviolet (UV) light.

Unless otherwise specified, negative control (-) sample was processed identically to experimental samples, except that buffer was added instead of ADP-ribosyltransferases during ADP-ribosylation step, providing an internal reference for unmodified oligonucleotide. Positive control (+) samples were also treated identically to experimental samples, except that buffer was added in place of hydrolases and other treatment conditions as described above during the de-modification step, serving as a reference for modified oligonucleotides.

## Acknowledgments

The use of the facilities and expertise of the Biocenter Oulu Structural Biology core facility (a member of Biocenter Finland, Instruct-ERIC Centre Finland and FINStruct), Proteomics and Protein Analysis core facility (a member of Biocenter Finland) and Biocenter Oulu sequencing center are gratefully acknowledged.

## Author contributions

M.T.H.D., L.L. conceptualization; M.T.H.D., Y.L. investigation; M.T.H.D. writing-original draft; M.T.H.D, Y.L, I.A., L.L. writing-reviewing and editing; I.A., L.L. funding acquisition; I.A., L.L. supervision.

## Abbreviations

CFP: mCerulean
DarG: DNA ADP-ribosylglycohydrolase
DarT: DNA ADP-ribosyltransferase
FRET: fluorescence resonance energy transfer
GAP: Gαi protein-based peptide
GdnHCl: Guanidine HCl
HTS: high-throughput screening
MARylated: mono-ADP-ribosylated
MD: macrodomain
M. tuberculosis: Mycobacterium tuberculosis
*Mtb*DarG: DarG in *M. tuberculosis*
nanoDSF: nano differential scanning fluorimetry
PAR: poly-ADP-ribose; PARylated, poly-ADP- ribosylated
PARG: poly(ADP-ribose) glycohydrolase
PARP: poly(ADP-ribose) polymerases
rFRET: ratiometric FRET
TA: toxin-antitoxin
TEV: tobacco etch virus
T_m_: melting temperature
YFP: mCitrine

## References

1. Takamura-Enya T, Watanabe M, Totsuka Y, Kanazawa T, Matsushima-Hibiya Y, Koyama K, Sugimura T & Wakabayashi K (2001) Mono(ADP-ribosyl)ation of 2’-deoxyguanosine residue in DNA by an apoptosis-inducing protein, pierisin-1, from cabbage butterfly. Proc Natl Acad Sci U S A 98, 12414–12419.

2. Lüscher B, Ahel I, Altmeyer M, Ashworth A, Bai P, Chang P, Cohen M, Corda D, Dantzer F, Daugherty MD, Dawson TM, Dawson VL, Deindl S, Fehr AR, Feijs KLH, Filippov DV, Gagné J-P, Grimaldi G, Guettler S, Hoch NC, Hottiger MO, Korn P, Kraus WL, Ladurner A, Lehtiö L, Leung AKL, Lord CJ, Mangerich A, Matic I, Matthews J, Moldovan G-L, Moss J, Natoli G, Nielsen ML, Niepel M, Nolte F, Pascal J, Paschal BM, Pawłowski K, Poirier GG, Smith S, Timinszky G, Wang Z-Q, Yélamos J, Yu X, Zaja R & Ziegler M (2022) ADP-ribosyltransferases, an update on function and nomenclature. FEBS J 289, 7399–7410.

3. Suskiewicz MJ, Prokhorova E, Rack JGM & Ahel I (2023) ADP-ribosylation from molecular mechanisms to therapeutic implications. Cell 186, 4475–4495.

4. Hassa P O (2008) The diverse biological roles of mammalian PARPS, a small but powerful family of poly-ADP-ribose polymerases. Front Biosci 13, 3046.

5. Yélamos J, Schreiber V & Dantzer F (2008) Toward specific functions of poly(ADP-ribose) polymerase-2. Trends Mol Med 14, 169–178.

6. Hottiger MO, Hassa PO, Lüscher B, Schüler H & Koch-Nolte F (2010) Toward a unified nomenclature for mammalian ADP-ribosyltransferases. Trends Biochem Sci 35, 208–219.

7. D’Amours D, Desnoyers S, D’Silva I & Poirier GG (1999) Poly(ADP-ribosyl)ation reactions in the regulation of nuclear functions. Biochem J 342 ( **Pt 2**), 249–268.

8. Yamada M, Miwa M & Sugimura T (1971) Studies on poly (adenosine diphosphate-ribose). Arch Biochem Biophys 146, 579–586.

9. Juarez-Salinas H, Levi V, Jacobson EL & Jacobson MK (1982) Poly(ADP-ribose) has a branched structure in vivo. J Biol Chem 257, 607–609.

10. Minaga T & Kun E (1983) Probable helical conformation of poly(ADP-ribose). The effect of cations on spectral properties. J Biol Chem 258, 5726–5730.

11. Lee JS, Latimer LJP & Sibley JT (1989) Detection of unusual nucleic acids with monoclonal antibodies specific for triplexes, poly(ADP-ribose), DNA-RNA hybrids, and Z-DNA. Anal Biochem 178, 373–377.

12. Kanai Y, Matsushima T & Sugimura T (1978) Induction of specific antibodies to poly(ADP– ribose) in rabbits by double-stranded RNA, poly(A)·poly(U). Nature 274, 809–812.

13. Schreiber V, Dantzer F, Ame J-C & de Murcia G (2006) Poly(ADP-ribose): novel functions for an old molecule. Nat Rev Mol Cell Biol 7, 517–528.

14. Gibson BA & Kraus WL (2012) New insights into the molecular and cellular functions of poly(ADP-ribose) and PARPs. Nat Rev Mol Cell Biol 13, 411–424.

15. Virág L, Robaszkiewicz A, Rodriguez-Vargas JM & Oliver FJ (2013) Poly(ADP-ribose) signaling in cell death. Mol Aspects Med 34, 1153–1167.

16. Feijs KLH, Forst AH, Verheugd P & Lüscher B (2013) Macrodomain-containing proteins: regulating new intracellular functions of mono(ADP-ribosyl)ation. Nat Rev Mol Cell Biol 14, 443– 451.

17. Rack JGM, Perina D & Ahel I (2016) Macrodomains: Structure, Function, Evolution, and Catalytic Activities. Annu Rev Biochem 85, 431–454.

18. Palazzo L, Mikolčević P, Mikoč A & Ahel I (2019) ADP-ribosylation signalling and human disease. Open Biol 9, 190041.

19. Leung AKL, McPherson RL & Griffin DE (2018) Macrodomain ADP-ribosylhydrolase and the pathogenesis of infectious diseases. PLOS Pathog 14, e1006864.

20. Mikolčević P, Hloušek-Kasun A, Ahel I & Mikoč A (2021) ADP-ribosylation systems in bacteria and viruses. Comput Struct Biotechnol J 19, 2366–2383.

21. Catara G, Corteggio A, Valente C, Grimaldi G & Palazzo L (2019) Targeting ADP-ribosylation as an antimicrobial strategy. Biochem Pharmacol 167, 13–26.

22. Page R & Peti W (2016) Toxin-antitoxin systems in bacterial growth arrest and persistence. Nat Chem Biol 12, 208–214.

23. Ramage HR, Connolly LE & Cox JS (2009) Comprehensive Functional Analysis of Mycobacterium tuberculosis Toxin-Antitoxin Systems: Implications for Pathogenesis, Stress Responses, and Evolution. PLoS Genet 5, e1000767.

24. Schuller M, Butler RE, Ariza A, Tromans-Coia C, Jankevicius G, Claridge TDW, Kendall SL, Goh S, Stewart GR & Ahel I (2021) Molecular basis for DarT ADP-ribosylation of a DNA base. Nature 596, 597–602.

25. Jankevicius G, Ariza A, Ahel M & Ahel I (2016) The Toxin-Antitoxin System DarTG Catalyzes Reversible ADP-Ribosylation of DNA. Mol Cell 64, 1109–1116.

26. Catara G, Caggiano R & Palazzo L (2023) The DarT/DarG Toxin–Antitoxin ADP-Ribosylation System as a Novel Target for a Rational Design of Innovative Antimicrobial Strategies. Pathogens 12, 240.

27. Deep A, Singh L, Kaur J, Velusamy M, Bhardwaj P, Singh R & Thakur KG (2023) Structural insights into DarT toxin neutralization by cognate DarG antitoxin: ssDNA mimicry by DarG C-terminal domain keeps the DarT toxin inhibited. Structure 31, 780–789.e4.

28. Butler RE, Schuller M, Jaiswal R, Mukhopadhyay J, Barber J, Hingley-Wilson S, Wasson E, Couto Alves A, Ahel I & Stewart GR (2025) Control of replication and gene expression by ADP-ribosylation of DNA in Mycobacterium tuberculosis. EMBO J 44, 3468–3491.

29. Tromans-Coia C, Sanchi A, Moeller GK, Timinszky G, Lopes M & Ahel I (2021) TARG1 protects against toxic DNA ADP-ribosylation. Nucleic Acids Res 49, 10477–10492.

30. Zaveri A, Wang R, Botella L, Sharma R, Zhu L, Wallach JB, Song N, Jansen RS, Rhee KY, Ehrt S & Schnappinger D (2020) Depletion of the DarG antitoxin in *Mycobacterium tuberculosis* triggers the DNA-damage response and leads to cell death. Mol Microbiol 114, 641–652.

31. Gygli SM, Borrell S, Trauner A & Gagneux S (2017) Antimicrobial resistance in Mycobacterium tuberculosis: mechanistic and evolutionary perspectives. FEMS Microbiol Rev 41, 354–373.

32. Nguyen L (2016) Antibiotic resistance mechanisms in M. tuberculosis: an update. Arch Toxicol 90, 1585–1604.

33. Floyd K, Glaziou P, Zumla A & Raviglione M (2018) The global tuberculosis epidemic and progress in care, prevention, and research: an overview in year 3 of the End TB era. Lancet Respir Med 6, 299–314.

34. Passador L & Iglewski W (1994) [49] ADP-ribosylating toxins. In Methods in Enzymology pp. 617–631. Elsevier.

35. Lu Y, Schuller M, Bullen NP, Mikolcevic P, Zonjic I, Raggiaschi R, Mikoc A, Whitney JC & Ahel I (2025) Discovery of reversing enzymes for RNA ADP-ribosylation reveals a possible defence module against toxic attack. Nucleic Acids Res 53, gkaf069.

36. Sowa ST, Galera-Prat A, Wazir S, Alanen HI, Maksimainen MM & Lehtiö L (2021) A molecular toolbox for ADP-ribosyl binding proteins. Cell Rep Methods 1, 100121.

37. Slade D, Dunstan MS, Barkauskaite E, Weston R, Lafite P, Dixon N, Ahel M, Leys D & Ahel I (2011) The structure and catalytic mechanism of a poly(ADP-ribose) glycohydrolase. Nature 477, 616–620.

38. Song Y, Rodgers VGJ, Schultz JS & Liao J (2012) Protein interaction affinity determination by quantitative FRET technology. Biotechnol Bioeng 109, 2875–2883.

39. Zhang J-H, Chung TDY & Oldenburg KR (1999) A Simple Statistical Parameter for Use in Evaluation and Validation of High Throughput Screening Assays. SLAS Discov 4, 67–73.

40. Shoichet BK (2006) Interpreting Steep Dose-Response Curves in Early Inhibitor Discovery. J Med Chem 49, 7274–7277.

41. Lalić J, Posavec Marjanović M, Palazzo L, Perina D, Sabljić I, Žaja R, Colby T, Pleše B, Halasz M, Jankevicius G, Bucca G, Ahel M, Matić I, Ćetković H, Luić M, Mikoč A & Ahel I (2016) Disruption of Macrodomain Protein SCO6735 Increases Antibiotic Production in Streptomyces coelicolor. J Biol Chem 291, 23175–23187.

42. Hloušek-Kasun A, Mikolčević P, Rack JGM, Tromans-Coia C, Schuller M, Jankevicius G, Matković M, Bertoša B, Ahel I & Mikoč A (2022) Streptomyces coelicolor macrodomain hydrolase SCO6735 cleaves thymidine-linked ADP-ribosylation of DNA. Comput Struct Biotechnol J 20, 4337–4350.

43. Glumoff T, Sowa ST & Lehtiö L (2022) Assay technologies facilitating drug discovery for ADP-ribosyl writers, readers and erasers. BioEssays 44, 2100240.

44. Keam SJ, Lyseng-Williamson KA & Goa KL (2003) Pranlukast: A Review of its Use in the Management of Asthma. Drugs 63, 991–1019.

45. Mishra A, Mamidi AS, Rajmani RS, Ray A, Roy R & Surolia A (2018) An allosteric inhibitor of *Mycobacterium tuberculosis* ArgJ: Implications to a novel combinatorial therapy. EMBO Mol Med 10, e8038.

46. Rajmani RS & Surolia A (2024) Antimycobacterial and healing effects of Pranlukast against MTB infection and pathogenesis in a preclinical mouse model of tuberculosis. Front Immunol 15, 1347045.

47. Jeong J-Y, Yim H-S, Ryu J-Y, Lee HS, Lee J-H, Seen D-S & Kang SG (2012) One-Step Sequence- and Ligation-Independent Cloning as a Rapid and Versatile Cloning Method for Functional Genomics Studies. Appl Environ Microbiol 78, 5440–5443.

48. Reyrat J-M, Pelicic V, Gicquel B & Rappuoli R (1998) Counterselectable Markers: Untapped Tools for Bacterial Genetics and Pathogenesis. Infect Immun 66, 4011–4017.

49. Hynes MF, Quandt J, O’Connell MP & Pühler A (1989) Direct selection for curing and deletion of Rhizobium plasmids using transposons carrying the Bacillus subtilis sacB gene. Gene 78, 111– 120.

50. Inglese J, Johnson RL, Simeonov A, Xia M, Zheng W, Austin CP & Auld DS (2007) High-throughput screening assays for the identification of chemical probes. Nat Chem Biol 3, 466–479.

