## Supplementary information for "A FRET-based high-throughput screening assay for the discovery of *Mycobacterium tuberculosis* DNA ADP-ribosylglycohydrolase DarG inhibitors"

### Content

**Figure S1**: FRET-based assay of CFP-*Mtb*DarG MD and MARylated YFP-GAP.

**Figure S2**: Effect of pH on the rFRET signals of CFP-*Mtb*DarG MD and PARylated YFP-GAP.

**Figure S3**: Effect of NaCl on the rFRET signals of CFP-*Mtb*DarG MD and PARylated YFP-GAP.

**Figure S4**: Effect of glycerol on rFRET signal stability.

**Figure S5**: Re-test the inhibition activity of 17 hit compounds on *Mtb*DarG MD and PARylated YFP binding interaction.

**Figure S6**: Scattering signals of *Mtb*DarG in the absence and presence of hits.

**Table 1**: Statistical parameters of assay validation.

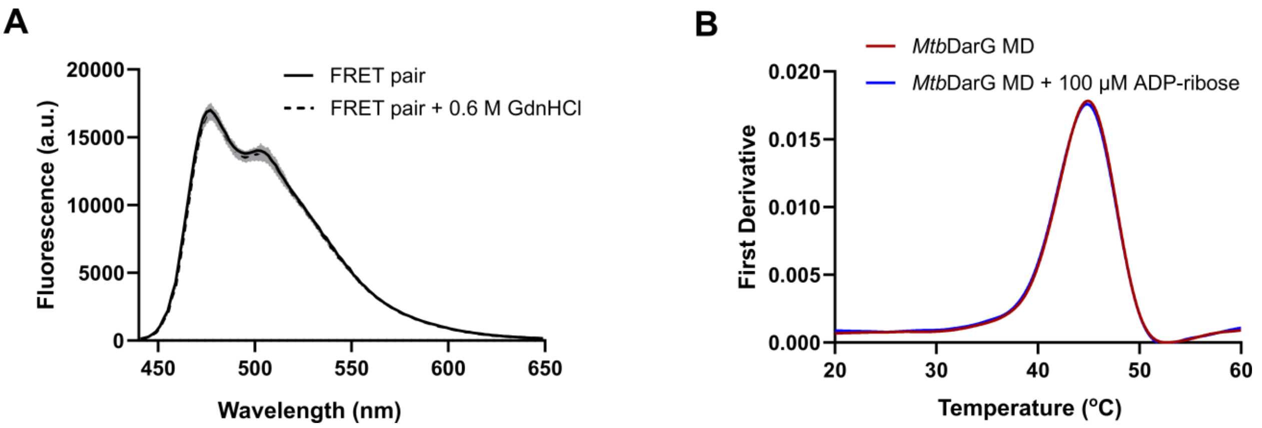


**Figure S1: (**A) The fluorescence spectra of 100 nM CFP-*Mtb*DarG MD and 500 nM MARylated YFP-GAP mixture in the absence (solid line) or presence (dashed line) of 0.6 M GdnHCl. The curves shown are mean ± SD of four replicates. (B) ADP-ribose did not stabilize *Mtb*DarG shown in nanoDSF.

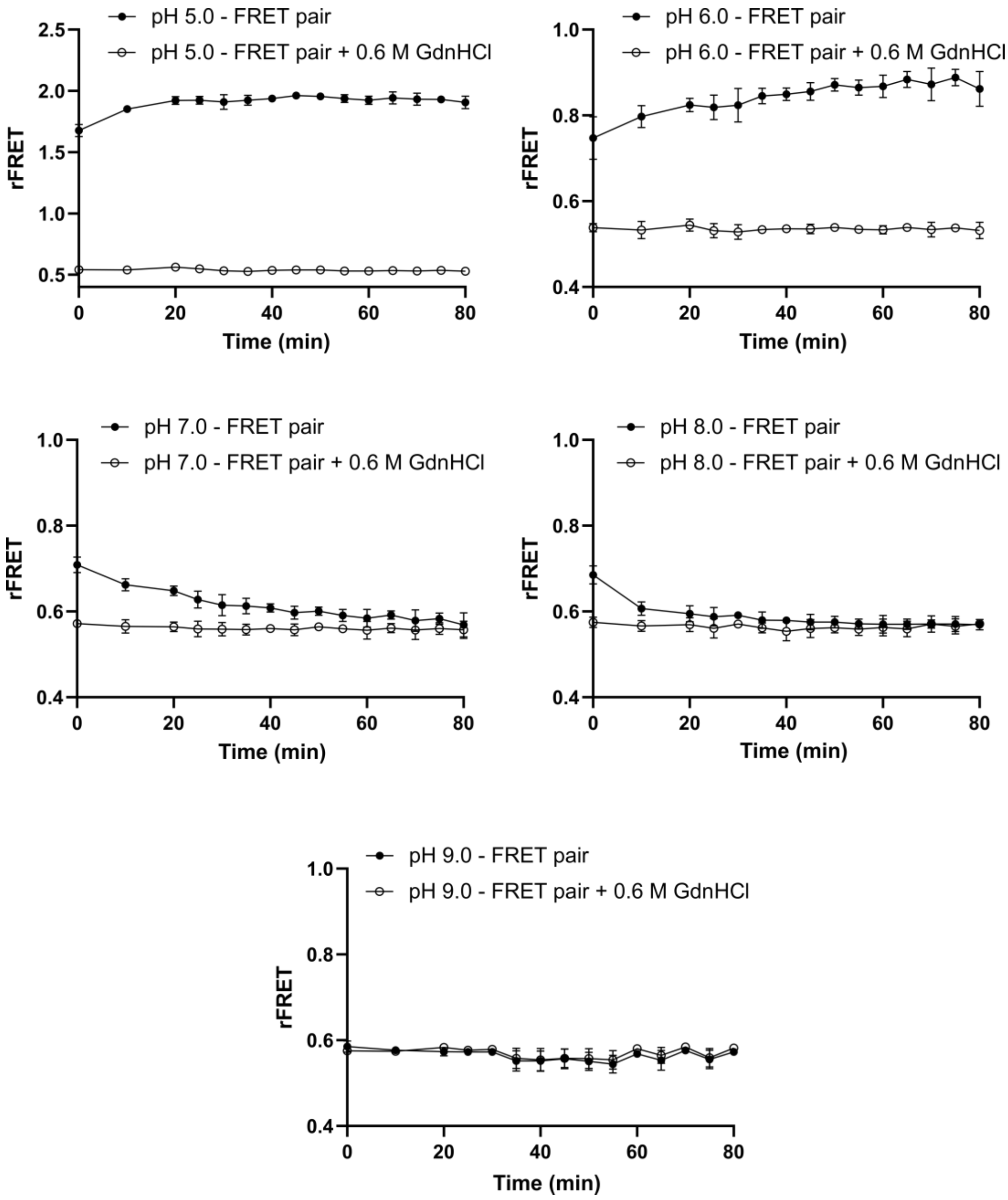


**Figure S2:** Effect of pH on the rFRET signal. 100 nM CFP-*Mtb*DarG MD was mixed with 500 nM PARylated YFP-GAP in different pH buffers. Buffer condition: 25 mM NaCl, 0.01% (v/v) Triton X-100, 0.5 mM TCEP in the presence of 10 mM MES pH 6.0, or 10 mM BTP pH 7.0, or 10 mM BTP pH 8.0, or 10 mM BTP pH 9.0. The signals were monitored for 80 min. Data shown are mean ± SD from 4 replicates.

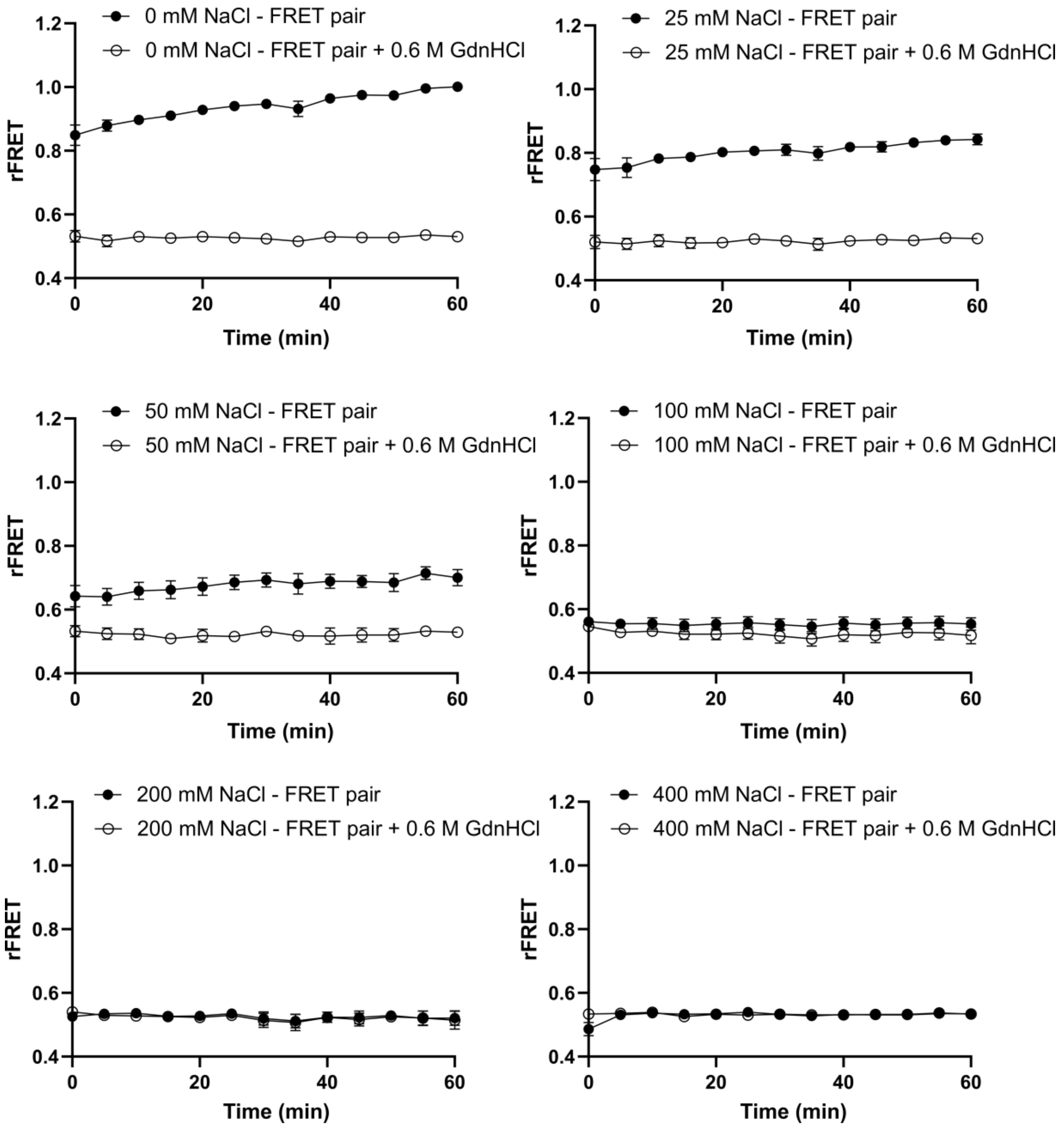


**Figure S3:** Effect of different NaCl concentrations on the rFRET signal. 100 nM CFP-*Mtb*DarG MD was mixed with 500 nM PARylated YFP-GAP in buffers containing different NaCl concentrations. Buffer included 10 mM MES pH 6.0, 0.01%(v/v) Triton X-100, 0.5 mM TCEP, 0 – 400 mM NaCl. The signals were measured during 60 min. Data shown are mean ± SD from 4 replicates.

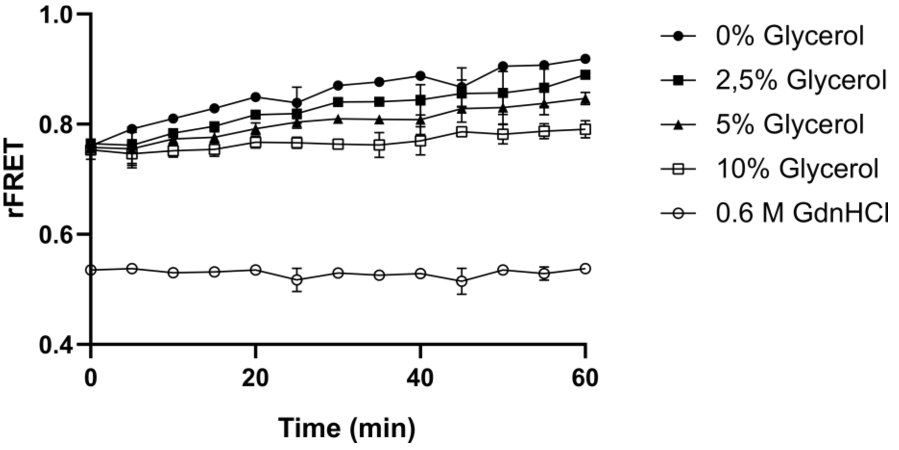


**Figure S4:** Effect of glycerol on rFRET signals. 100 nM CFP-*Mtb*DarG MD was mixed with 500 nM PARylated YFP-GAP in buffer containing 10 mM MES pH 6.0, 25 mM NaCl, 0.01% (v/v) Triton X-100, 0.5 mM TCEP, 0-10% (v/v) glycerol. The stability of signals was measured during 60 min. Data shown are mean ± SD from 4 replicates.

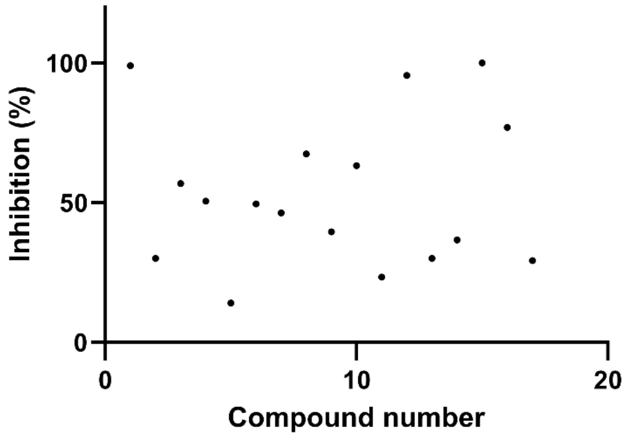


**Figure S5**. Seventeen hit compounds obtained from initial screening were re-tested using *Mtb*DarG MD-PARylated GAP FRET-based assay. All compounds showed inhibitory activity against the binding interaction of CFP-*Mtb*DarG MD and PARylated YFP-GAP. Data shown are mean of duplication.

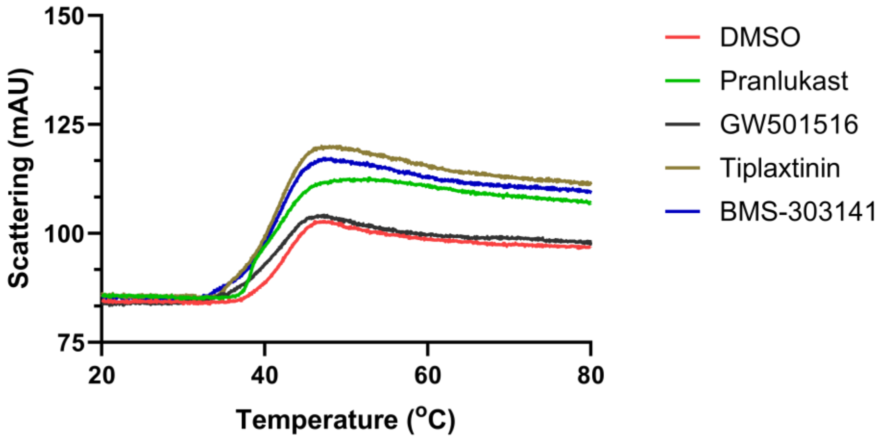


**Figure S6**. Scattering signal of *Mtb*DarG MD in the absence or presence of four hits. 0.3 mg/ml of *Mtb*DarG was incubated with 60 μM of compounds. Three measurements were taken. One presentive curve for each sample was shown.

**Table S1**: The statistical parameters of assay validation.

| Parameters | Day 1 | Day 2 | Day 3 | | |
|---|---|---|---|---|---|
| | | | Plate 1 | Plate 2 | Plate 3 |
| Positive control AVR (rFRET) ± SD (CV%) | 1.03 ± 0.02 (2.02%) | 0.97 ± 0.02 (1.62%) | 0.92 ± 0.02 (1.71%) | 0.96 ± 0.01 (1.37%) | 0.94 ± 0.01 (1.18%) |
| Negative control AVR (rFRET) ± SD (CV%) | 0.76 ± 0.01 (1.08%) | 0.73 ± 0.01 (1.35%) | 0.71 ± 0.01 (0.81%) | 0.74 ± 0.01 (0.82%) | 0.73 ± 0.01 (0.96%) |
| Z’-factor (CV%) | 0.68 | 0.68 | 0.68 | 0.74 | 0.75 |
| | 0.71 ± 0.04 (5.00%) | | | | |
| S/B (CV%) | 1.36 | 1.33 | 1.29 | 1.30 | 1.29 |
| | 1.31 ± 0.03 (2.20%) | | | | |
| S/N (CV%) | 32.86 | 24.56 | 35.40 | 36.95 | 32.05 |
| | 32.36 ± 4.78 (14.78%) | | | | |
| Plate-to-plate CV (%) Calculated from Z’-factor | 4.91% | | | | |
| Day-to-day CV (%) Calculated from Z’-factor | 3.77% | | | | |
